# Rewarding Value or Prediction Error: Settling the debate over the role of dopamine in reward learning

**DOI:** 10.1101/2022.11.06.515338

**Authors:** Alexandra A. Usypchuk, Etienne JP Maes, Megan Lozzi, Matthew P.H. Gardner, Geoffrey Schoenbaum, Guillem R. Esber, Mihaela D. Iordanova

## Abstract

The discovery that DA transients can be mapped onto the reward prediction errors in temporal difference models is a pinnacle achievement of neuroscience. Yet, there is abundant evidence that DA activity reinforces actions, suggesting it serves as an intrinsically rewarding event. These two possibilities are so conceptually intertwined that it is not surprising that they have been so far experimentally conflated. Here, using computational modeling, behavioural blocking and optogenetics, we show that stimulating VTA DA neurons promotes learning even when a natural reward and DA stimulation are held constant across the learning phases of blocking. These findings provide strong evidence in favour of the prediction error hypothesis rather than encoding the rewarding value of appetitive events.

---

It is incontrovertible that VTA DA stimulation acts as a reinforcing signal for actions and states (e.g., Olds & Milner, 1954; Wise, 1978; Schultz et al., 1997). Animals readily self-stimulate for, and frequent places where, electrical or optogenetic activation of VTA DA neurons occurred, and disrupting this activity prevents learning or reduces established reward-seeking responses (e.g., Carter et al., 2022; Corbett & Wise, 1980; Crow, 1972; Ilango et al., 2014; Pascoli et al., 2015; Phillips & Fibiger, 1973; Millard et al., 2022; Witten et al., 2010). An intuitive interpretation of these findings is that VTA DA activity constitutes an appetitive event that is intrinsically rewarding. An alternative is that VTA DA activity encodes a reward prediction error - the difference between the predicted and obtained reward – which provides a teaching signal that drives learning (Schultz et al., 1997). These two possibilities are so conceptually intertwined that it is not surprising that they have been so far experimentally conflated. For example, on the one hand changes in value provided by the VTA DA signal would necessarily modulate prediction-error and therefore learning. Indeed, studies reporting new learning by optogenetically increasing DAergic activity could be doing so by boosting the rewarding value of the event present during the stimulation either and thus generating positive prediction errors to drive learning. Similarly, studies reporting behavioral extinction after optogenetically suppressing DAergic activity might be achieving this effect either by blunting the rewarding value of the event upon which the inhibition occurs and thus creating negative prediction errors. An alternative holds that rewarding value and prediction error are two dissociable processes, which are capable of being decorrelated. For example, while the presence of a prediction error will always drive learning, the presence of a reward signal may support reward-seeking behaviors without necessarily inducing learning (e.g., when the reward is expected).

Given that DA is at the heart of reward processing and reward-related disorders, it is fundamental that we understand the relationship between rewarding value and prediction error, as this would determine the mechanisms of adaptive and maladaptive reward pursuit. To dissociate these alternatives, we combined computational modeling, behavioural blocking and optogenetics. In blocking, learning about a cue➔reward association is attenuated in the presence of a previously established predictor for the same reward. This association will form, however, if the magnitude of reward is increased across learning phases (unblocking; Mackintosh, 1977; Dickinson, 1976) or VTA DA neurons are stimulated at the time of the *expected* reward (Steinberg et al., 2013; Keiflin et al., 2019). Based on these findings, we reasoned that if VTA DA neurons signal the presence of a rewarding event, then delivering stimulation during the conditioning phase, so that it can become expected just like any other reward, should prevent unblocking when it is delivered later when a second cue is added. Conversely, if VTA DA neurons encode reward prediction errors then by definition it cannot become expected, and stimulation in the second phase should result in unblocking, regardless whether the identical stimulation had been delivered with the reward during conditioning.

## Optogenetic stimulation of VTA DA transients promotes learning in blocking

Before settling whether VTA DA transients produce a rewarding event or a prediction error, we confirmed, computationally and experimentally, that optogenetic stimulation of VTA DA neurons at the time of expected reward delivery in a blocking design drive learning (unblocking). Fifteen Long Evans transgenic rats expressing Cre recombinase under the control of the tyrosine hydroxylase promoter (TH-Cre^+/-^) received viral infusion (AAV5-EF1α–DIO-ChR2–eYFP) and optical fiber implantation bilaterally in the medial VTA. Rats were food-restricted for two days and then trained using a within-subjects blocking design (Fig 1A), which consisted of conditioning, blocking and test. During conditioning, the rats learned an A➔food and B➔nothing discrimination, evident in greater percent time spent in the food cup during A compared to B (Supp Fig 1A, F(1,14) = 26.054, p<0.001, n^2^ = 0.650). Subsequently, during blocking, A was presented in compound with either X or Y to create two reinforced blocking compounds, AX➔food and AY➔food, whereas B was presented in compound with Z to create a reinforced control compound, BZ➔food. Optical stimulation of VTA TH+ neurons occurred during reward delivery on AX➔food trial and during the intertrial interval following AY➔food trials. There were no differences in responding among the three compounds by the end of the blocking phase (Supp Fig 1A; min *p* = .087, 95% CI [−.393, 7.117]). Our data (Fig 1C: Test) show blocking to Y (i.e., low levels of responding, Y vs. Z: *p* = .023, 95% CI [−1.333, −19.307]) and unblocking to X (i.e., high levels of responding, X vs. Y: *p* = .022, 95% CI [1.080, 14.942]). Further, the unblocking effect to X was complete (i.e., responding to X did not differ from Z, *p* = 1.00, 95% CI [−11.550, 6.932]). Computational modelling of VTA DA stimulation as a rewarding event (value) or a prediction error predicted learning about X (Fig 1D; Supp Fig 1B and 1C), which our data confirmed. While the results replicate and extend previous findings, they do not disentangle the two competing hypotheses.

**Figure 1.**
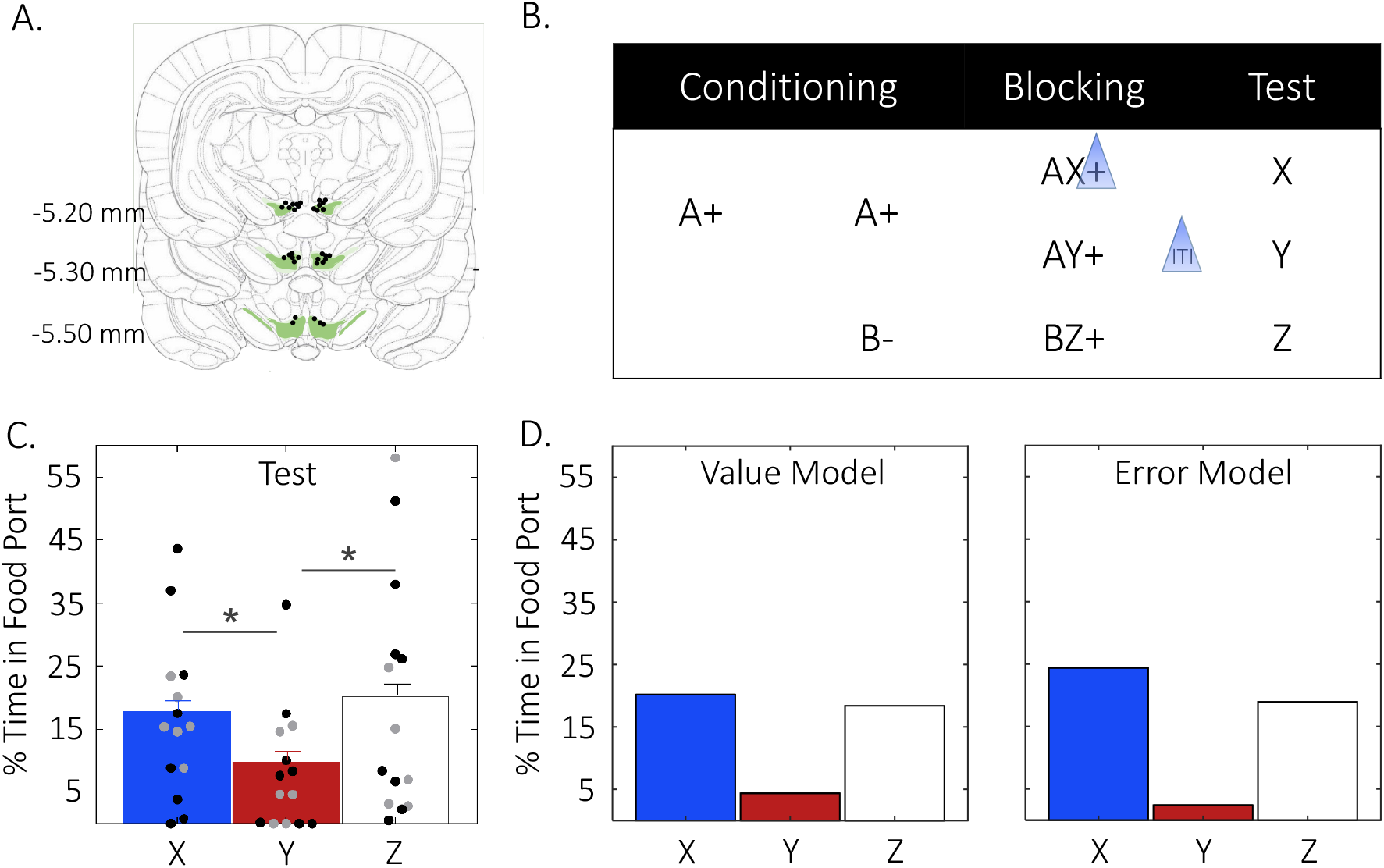
Optogenetic stimulation of VTA DA transients promotes learning in blocking. **A)** Optical fiber implant placements and virus expression in the VTA. **B**) Experimental design. Blue triangles indicate optical stimulation. **C**) Behavioral results (mean + s.e.m., *n* = 15 rats; males in gray, females in black) on test confirm an effect of cue (ANOVA: *F_Cue_*(2,28) = 6.070, *p* = .006, *η^2^* = .302) where responding to the blocked cue Y was lower than to the control cue Z (*M_diff_*= −10.320, *SE* = 3.307, *p* = .023, 95% CI [−1.333, −19.307) while responding to the unblocking cue X was greater than the blocked cue Y (*M_diff_* = 8.011, *SE* = 2.550, *p* = .022, 95% CI [1.080, 14.942) but no different than Z (*M_diff_*= −2.309, *SE* = 3.400, *p* = 1.00, 95% CI [−11.550, 6.932]). **D**) Predicted results based on computational modeling where VTA DA stimulation acts as a rewarding event (Value Model; left) or as a prediction error (Error Model; right). Note the output of the classic temporal difference reinforcement learning model was converted from *V* to CR to better reflect the behavioral output actually measured in our experiments (see Methods for details).

## Optogenetic stimulation of VTA DA transients generate a prediction error, not a rewarding event

To determine whether inducing VTA DA transients drives learning by generating a prediction error or producing a rewarding event, we optically stimulated VTA dopamine neurons at the time of food delivery during both conditioning and blocking (Fig 2a). If DA transients mimic a rewarding event, then holding the DA stimulation constant across the two phases should abolish the unblocking observed in the above experiment, when it was only delivered in the blocking phase. In contrast, if DA transients signal a prediction error, then unblocking should be unaffected by delivery in conditioning.

**Figure 2.**
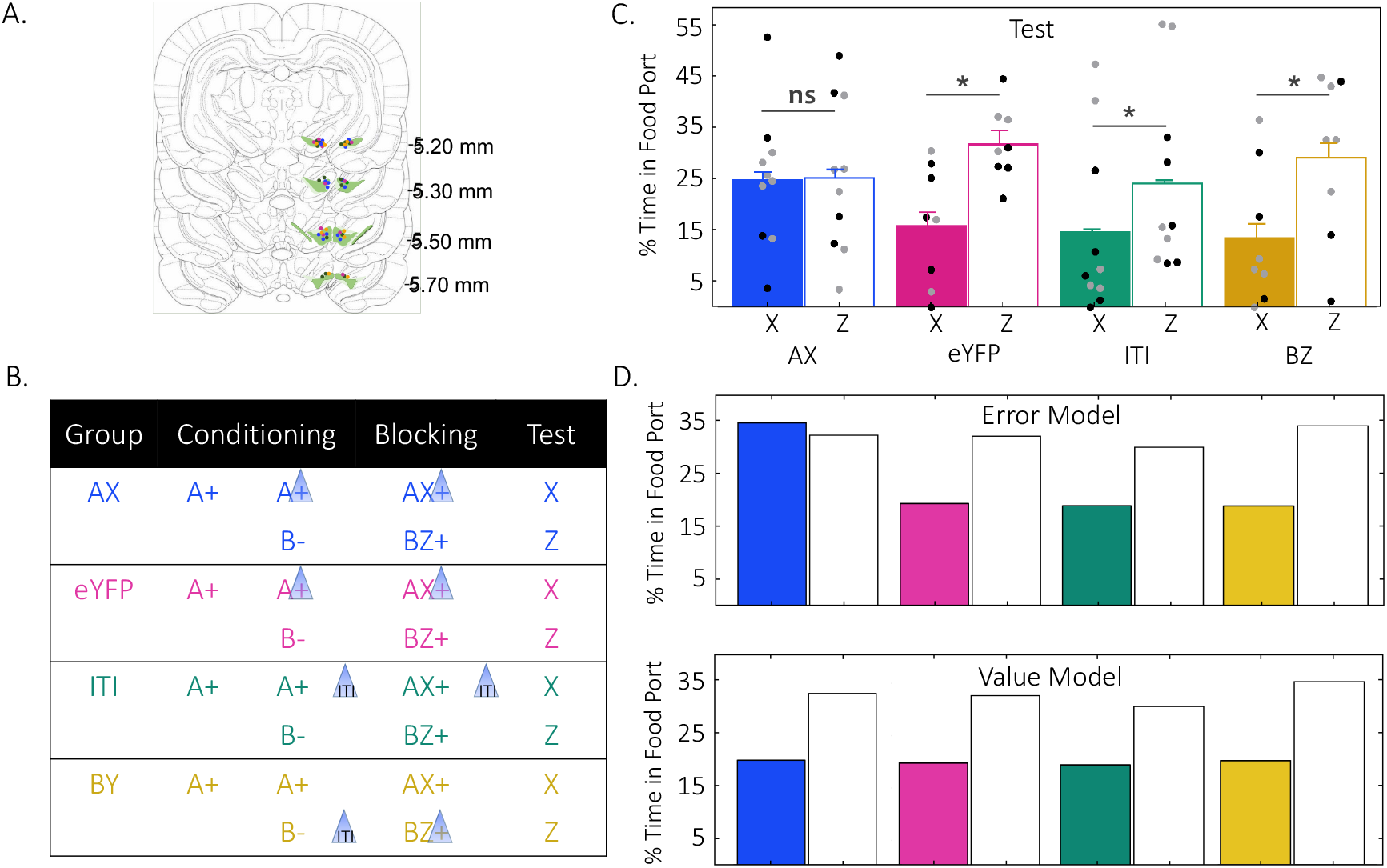
Optogenetic stimulation of VTA DA transients generate a prediction error, not a rewarding event. **A)** Optical fiber implant placements and virus expression in the VTA (blue = AX, fuchsia = eYFP, green = ITI, gold = BZ) **B)** Behavioral design. Blue triangles indicate optical stimulation. **C)** Behavioral results (mean + s.e.m., *n* = 44 rats; males in gray, females in black) on test confirm a cue by group interaction (ANOVA: *F_Cue x Group_*(3,32) = 3.757, *p* = .020, *η^2^* = .260). Responding to X was lower than to the control cue Z for all groups (max *M_diff_* = −10.000, *SE* = 4.040, *p* = .019, 95% CI −18.230, −1.771]) except Group AX (*M_diff_* = - .430, *SE* = 4.062, *p* = .916, 95% CI [−8.704, 7.844]). **D)** Predicted results based on computational modeling where VTA DA stimulation acts as a reward prediction error (Error Model; top) or a rewarding event (Value Model; bottom). Note the output of the classic temporal difference reinforcement learning model was converted from *V*to CR to better reflect the behavioral output actually measured in our experiments (see Methods for details).

Thirty-six TH-Cre^+/-^ rats received viral infusion (AAV5-EF1α–DIO-ChR2–eYFP or AAV5-EF1α–DIO–eYFP) and bilateral optical fiber implantation in the medial VTA (Fig 2A). Each of four groups were trained in a within-subject blocking design (Fig 2B) similar to the one described earlier, consisting of a blocked cue (X) and a control cue (Z). Optical stimulation of TH+ neurons differed among the four groups. The critical experimental condition, Group AX, received stimulation during reward delivery on A→food (conditioning) and AX→food (blocking) trials. To control for light artifacts, Group eYFP (lacking the opsin in TH+ neurons) received identical optical stimulation to Group AX. To control for the temporal specificity of stimulation, Group ITI was stimulated during the ITI in both phases of learning (conditioning, blocking). To further test if stimulation of VTA TH+ neurons induced a rewarding event additional to the natural reward delivered (food), a third control was generated, Group BY, which received stimulation during the ITI following B➔nothing trials in conditioning and during reward delivery on control trials in blocking (i.e., BZ→ reward).

Rats learned the A➔food and B➔nothing discrimination (Supp Fig 2A, *F* (5,160) = 4.328, *p* = .001, *η^2^* = .119) during conditioning, and by the end of blocking each group responded similarly to AX and BZ (Supp Fig 2A; min *p* = .103, CI [-2.033, 21.214]). Responding on test revealed unblocking in Group AX (Fig 2C; similar responding to X and Z, *p* = .916, 95% CI [-8.704, 7.844]) and blocking in all other groups (i.e., responding lower to X compared with Z, max *p* = .019, [−18.230, −1.771]). This pattern of results is consistent with the computational model that envisages VTA DA transients as a reward prediction error and inconsistent with the proposal that it serves as a rewarding event (value, Fig 2D, Supp Fig 2B and 2C).

## Discussion

Here we have shown that when optical stimulation of VTA DA neurons occurs at the time of expected reward delivery (i.e., blocking phase), unblocking is obtained, in line with prior reports (Steinberg et al., 2013, Keiflin et al., 20XX). Remarkably, using the same stimulation in each of the two learning phases of the blocking design, and thus holding the putative level of reward constant across phases, did not disrupt the unblocking effect of DA stimulation. As illustrated by our computational modeling, this result is as expected if VTA DA stimulation is acting as a prediction error and cannot be explained if instead it acts as a rewarding event.

This conclusion is further bolstered by our finding that VTA DA stimulation does not boost the value of the natural reward utilized in our study, evident in the lack of difference in responding to Z on test in the BZ group relative to the other groups. Moreover, an increase in value with VTA DA stimulation was unlikely due to inadequate stimulation levels – unblocking was obtained in both experiments with this stimulation (responding to X was similar to that of Z in the experimental groups).

Although our data favours the reward prediction error hypothesis, it is well-known that different populations of VTA DA neurons subserve different functions (Cohen et al., 2012; Engelhard et al., 2019; Howe et al., 2016; Schultz et al., 1997). Therefore, we must allow for the possibility that distinct subpopulations of DA neurons might signal rewarding events and reward prediction errors. Here, indiscriminately driving VTA DA neurons encouraged learning by shifting the system towards inducing a reward prediction error. The dominance of each population, however, may be determined by the context in which it is tested. For example, stimulation in the context of classical conditioning drives a prediction error signal (e.g., Steinberg et al., 2013; Keiflin et al., 2018; Sharpe et al., 2017), whereas stimulation in an instrumental setting delivers rewarding value (e.g, Carter et al., 2022; Witten et al., 2010). While there may be some rewarding property embedded in the general DA signal, an error mechanism would parsimoniously account for the reinforcement property of VTA DA activation without having to appeal to the induction of rewarding value.

To our knowledge, ours is the first demonstration to explicitly pit the error vs. reinforcing value accounts of DA function against one another under otherwise identical conditions. Our data settle this issue and join a series of elegant studies that show prediction error-like signals during learning about valueless sensory events (Sadacca et al., 2016; Takahashi et al., 2017).

## METHODS

### Subjects

Transgenic rats expressing Cre recombinase under the control of the tyrosine hydroxylase (TH) promoter on a Long Evans background (TH:Cre rats) were used in the present experiments. Following histological analyses, the final number of rats included in the analyses was 15 (7 males, 8 females; weight 250-350g) and 36 (18 males, 18 females; weight 250-420g) for the first and second experiment, respectively. For source, housing, feeding and maintenance details see supplemental materials. All experimental procedures were in accordance with the Canadian Council on Animal Care and the Concordia University Animal Care Committee.

### Surgery

Rats were anaesthetized with isoflurane gas and placed on the stereotaxic apparatus (David Kopf Instruments) where they received bilateral VTA viral-infusions (AAV5-EF1α–DIO-ChR2–eYFP or AAV5-EF1α–DIO-eYFP) at the following coordinates relative to bregma: AP: −5.3mm; ML: 0.7mm; DV: −6.5mm and −7.7 (females) or −7.0mm and −8.2mm (males). At each site, 1 μL of virus was infused at a rate of .1 μL/min (10 min infusion; 10 min diffusion) through a blunt injector. Polyethylene tubing (Scientific Commodities Inc., Lake Havasu City, AZ) filled with dH_2_O connected the injector to a 10 μL syringe (Hamilton, NV) mounted on a microinfusion pump (Harvard Apparatus, MA). Optical fibers were implanted at the following coordinates relative to bregma: AP: −5.3mm; ML: ±2.61mm, and DV: −7.05mm (female) or −7.55mm (male) at an angle of 15° pointed toward the midline, and secured using six screws and dental acrylic. Subcutaneous injections of saline (2x 5ml/kg) and rimadyl solution (5 mg/kg; Pfizer, Kirkland, QC) were administered during surgery to prevent dehydration, as well as reduce inflammation and pain. Rats were monitored post-operatively and oral cephalexin was administered (50mg/ml; TEVA, Toronto, ON) for seven days. Behavioural training began between two and three weeks following viral infusion to ensure four weeks of neuronal transfection at the start of stimulation.

### Apparatus

#### Behavioral Chambers

Behavioural procedures were conducted in eight operant chambers (Med Associates, St. Albans, VT, USA) enclosed in melamine ventilated cabinets. Cameras were mounted on the back wall of the cabinet which fed to a monitor in an adjacent room for behavioural observation. Each cabinet measured 31.8 cm in height x 26.7 cm in length x 25.4 cm in width. The floors were made of stainless-steel rods, 4 mm in diameter and spaced 15 mm apart, with a tray below. The front door, back wall, and ceiling were made of clear Perspex and the left and right modular walls were made of aluminum. A red house light mounted 1cm below the ceiling and located on the back wall of the melamine boxes was used to illuminate the chamber. Panel lights used to deliver visual stimuli were mounted to the left (4-Hz flashing light) and right (steady light) side of the wall 15 cm above the floor. Auditory stimuli were delivered through a loud-speaker located at the front of the behavioural chamber on the floor of the melamine cabinet. The intensity of auditory stimuli were calibrated using a digital sound level meter (Tenma, 72-942). A recessed magazine was located on the center panel of the righthand modular wall, 3.9 cm above the floor. Pellet dispensers used for sucrose delivery were located on the right-hand wall outside the behavioural chambers, and were connected to the magazine inside. All stimuli were controlled by Med Associates software on the computer in the adjacent room.

#### Stimuli

Pellet dispensers delivered two 45-mg chocolate-flavored sucrose pellets (Bio-Serv, Flemington, NJ, USA) into a recessed magazine. The white noise (72 dB), clicker (71 dB), and buzzer (77-78 dB) were used in experiment 1 while white noise (72 dB) and siren (74 dB) were used in experiments 2 and 3. The background noise was 48-50 dB and 59-60 dB when the laser power generator was off and on, respectively. All stimuli were counterbalanced.

#### Optogenetic Equipment

Each box was equipped with a 150 mW, 473 nm laser and patch cord (Shanghai Laser & Optics Century Co.) that connected to a bilateral optical rotary joint (Doric Lenses, QC). Two custom-made patch cords 40-41 cm long delivered light from the rotary joint to the custom-made optical fiber implants (see Supplemental Material for details). Light pulses were delivered at 20 Hz (5 ms pulses) in a 2 s train starting at 9 s in Experiment 1 and 11s in Experiment 2. Prior to each session, laser output was measured using a power meter (ThorLabs, NJ) and verified by an oscilloscope (Tektronix TDS 2014) and set at 20 mW at the tip of the patch cords.

##### Patch cords

Two ceramic ferrules (diameter = 230 μm; equip #FG200UEa, ThorLabs) were attached to an optic fiber (diameter = 200 μm; FG200UEA, ThorLabs) using heat-dry epoxy. The patch cords were reinforced with furcation tubing (equip #FT020, ThorLabs) covered by metal cabling (equip #FT05SS, ThorLabs). Patch cords were connected to optical implants via blacked out 2.5 mm ceramic split sleeves (F1-8300SSC-25, Fiber Optics for Sale Co.)

##### Optical fiber implants

Heat-dry epoxy was used to secure an optical fiber (200 μm diameter; FG200UEA, ThorLabs) in stainless alloy ferrules of 2.5 mm length and 230 μm bore (F1-0064F-25, Fiber Optics for Sale Co.). Optical fiber implants were tested to have a uniform circular light output and minimum 90% laser light transmittance. Bilateral implants were matched based on percent transmittance.

### General Behavioral Procedures

All subjects were handled by the experimenter for five days prior to the start of the experiment. Behavioral training was conducted during the dark cycle. Sessions began with a 2 min adaptation period. Each reinforced conditioning trial consisted of a 10s cue (or compound) followed by pellet delivery at 9 s and 10 s. Non-reinforced trials consisted of cue alone presentations. The intertrial interval lasted on average 6 min (range: 4-8 min). Rats remained in the conditioning chamber for 2 min following the final trial. The walls, ceiling, floor and stainless-steel tray were cleaned with a 10 % acetic acid solution before each session.

#### Magazine Training

Rats were exposed to sucrose pellets (20 pellets/rat) in their home cages 24 h prior to magazine training. Each magazine training session lasted 40 min during which one sucrose pellet was delivered into the food port every 60 s.

#### Conditioning

Conditioning took place across 12 days. On days 1-6, rats received 16 presentations of a 10 s stimulus (A) paired with two sucrose pellets. On days 7-12, six non-reinforced presentations of visual cue (B) were intermixed with the 16 reinforced trials of A in two blocks such that B was presented randomly three times for every eight presentations of A. For the first experiment (data presented on Figure 1), the rats were tethered to the optical fiber cables starting on day 7 while optical stimulation began on day 13 (first day of blocking). For the second experiment (data presented in Figure 2), the rats were tethered to the optical fiber cables starting on day 0 and stimulation began on day 7. In both optogenetic experiments, rats were habituated to the background noise from the laser equipment one day prior to use.

#### Blocking

Each compound conditioning session began with eight reminder trials of A and B presented in the following order A A B B A A B B and counterbalanced for initial cue presentation across days. Following reminder trials, compounds were presented six times each in a pseudorandomized order. For the first experiment, this order was AX AX AY AY BZ BZ BZ BZ AY AY AX AX AY AY BZ BZ AX AX. For the second experiment the order was AX AZ AZ AX AZ AX AX AZ AX AZ AX AZ for day 1 and 3 and AX AZ AX AZ AX AX AZ AX AZ AZ AX AZ for days 2 and 4.

#### Test

To determine the level of conditioned approach acquired by the auditory stimuli, rats received three non-reinforced presentations of each cue. In the first experiment the order was: X Y Z Z X Y Y Z X, with two reinforced presentations of A after three consecutive non-reinforced trials. In the second experiment the order was X Y Y X Y X Y. Cues were counterbalanced for the initial stimulus presentation.

### Histology

To confirm viral expression and fiber optic placement where applicable, rats were euthanized with 1ml of a sodium pentobarbital solution diluted 1:1 with sodium chloride and immediately perfused with phosphate buffered saline (PBS) followed by 4 % paraformaldehyde solution. Brains remained in post-fix 4 % paraformaldehyde solution for 24 h followed by 20 % sucrose solution for 48 h. Brains were sectioned at 40 μm and virus expression and optical fiber implant placements were determined using the boundaries defined in the rat brain atlas (Paxinos & Watson, 1997).

### Quantification and Statistical Analysis

The percent time animals spent in the magazine during stimulus presentation was used as the conditioned response measure. Data were analyzed in SPSS 24.0 (IBM, New York, USA) using analysis of variance (ANOVA) with a significance level set at α = 0.05. Pairwise comparisons were assessed with standardized confidence intervals (CIs; 95% for the mean difference) and measures of effect size (ηp2 for ANOVA and Cohen’s d for pairwise comparisons; see Cohen, 1988) and reported with the Bonferroni correction.

### Modelling

Simulations of the behavioral designs were run using a one-step temporal difference learning algorithm, TD(0)14. This algorithm was used to estimate the value of different states of the behavioral paradigm with states being determined by the stimuli present at any particular time. Linear function approximation was used to estimate the value, *V*, of a given state, *s_t_*, by the features present during that state according to:

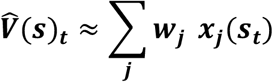

where *j* is indexed through all possible components of the feature vector **x** and corresponding weight vector **w**. The feature vector is considered to be the set of possible observed stimuli such that, if stimulus *j* is present during state *s* at time *t*, then **x***_j_*(*s_t_*) = 1, and 0 otherwise. The weights are adjusted over time to give the best approximation of the value of each state given the current set of stimuli. Weights, **w***_j_*, corresponding to each feature, **x***_j_*, are updated at each time step according to the temporal difference error rule:

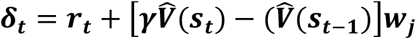

under linear value function approximation where *γ* is the temporal discounting factor. The weights are updated as:

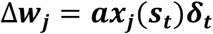

in which the scalar α is the learning rate. The linear value approximation reduces the size of the possible state space by generalizing states based on the features present. This approximation results in the calculation of the total expected value of a state as the sum of the expected value of each stimulus element present in the current state, a computation that is consistent with a global prediction error as stipulated by the Rescorla–Wagner model15.

### Model 1: Dopamine transients correspond to error

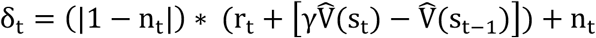

Where η is a value bounded between 0 and 1. This model was designed to capture the phenomena that as optogenetic stimulation increases, putatively more DA neurons will be recruited and activated, with complete saturation at η = 1. As such, any endogenously computed error signal will be swamped out in the condition η = 1, and more generally, this model assumes that mitigation of the endogenous error signal scales linearly with optogenetic stimulation and recruitment of DA neurons.

### Model 2: Dopamine transients correspond to value signals

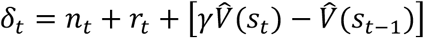

Where η is a value bounded between 0 and 1. Although dopamine stimulation acts as an additive term in both models, the comparator between the state values is unaffected in Model 2, and the adjusted state value, 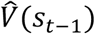, due to the stimulation term, *η*, can be estimated over time – *η* is not distinguishable from *r*. In Model 1, this is not the case. Here, stimulation replaces the comparator; such that with maximal stimulation the prediction error, δ_t_, is solely controlled by the stimulation, thus creating a condition in which the state value, 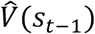, can not be estimated over time.

### Model parameterization

Generalization of value across stimuli was modeled by setting the initial weights, **w***j*, of a stimulus to 0.7 for stimuli of the same modality and 0.2 for stimuli of different modalities.

Conditioned responding to the food cup, CR, at each state was modeled using a logistic function:

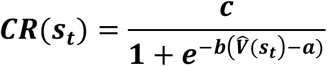

in which the parameters were determined based on empirical estimates of the maximal responding, *c*, the baseline responding, *a*, as well as the steepness of the learning curve, *b*. These were set as 55, 0.4 and 3, respectively, for all simulations. Reduced responding to the food cup while rats were attached to the patch cables was modeled as a reduction in the maximal responding to 40.

All simulations were performed with *α* = 0.05 and *γ* = 0.95. To ensure that the order of cue presentations did not affect the findings, cue presentations during each stage of conditioning were pseudo-randomized and the results of the simulations were averaged over 100 repetitions of the model. Simulations were performed using custom-written functions in MATLAB (Mathworks), which are available in the Supplementary Software and are posted on Github

## ACKNOWLEDGEMENTS

This work was supported by Natural Sciences and Engineering Research Council of Canada Discovery Grant (NSERC; RGPIN-2015-03658 to MDI); the Canada Research Chairs program (Grant # 950-230456 to MDI); a Graham Bell NSERC Graduate Scholarship (AAU); Fonds de Recherche du Quebec Nature et Technologie scholarship (AAU), a McGill University Hilton J McKeown Scholarship (EJPM), Concordia Undergraduate Student Research Award (MAL). All correspondence to be addressed to Mihaela D. Iordanova.

## AUTHOR CONTRIBUTIONS

AAU and MDI designed the experiments, interpreted the results, and wrote the manuscript. AAU, ML, EJPM performed the behavioral experiments, analyzed the data, and wrote the first draft of the manuscript. AAU, MDI and MG performed the computational modeling. GS provided animals for this project. GS and GER helped write the manuscript.

## CONFLICT OF INTEREST

The authors report no conflict of interest.

## Notes

### Competing Interest Statement

The authors have declared no competing interest.

