## Supplemental for "Rewarding Value or Prediction Error: Settling the debate over the role of dopamine in reward learning"

**Experiment 1: Optogenetic stimulation of VTA DA transients reinstates learning in blocking.**

Sex Effects. We used males and females in the studies reported. To used sex as a variable in all of our analyses to determine if males and females responded differently in our studies. With the exception of two instances (Experiment 1: Conditioning Days 1-6 returned a main effect of sex; Experiment 3: Blocking returned an interaction between sex and day), we found no effect of sex (max F) nor interactions with sex (max F) across training and test (Conditioning, Blocking, Test) in our studies. Therefore, we collapsed our data across males and females for all of our studies with the exception of the two instances mentioned above, the analyses of which are reported in the corresponding sections below. Table 1 contains all the statistical analyses pertaining to sex as a factor.

| **Experiment 1** $n_m = 7$; $n_f = 8$ | **Variable** | **df** | **F** | ***p*** | $\eta^2$ |
|---|---|---|---|---|---|
| Conditioning Day 1-6 | Sex | 1, 13 | 5.205 | .040* | .286 |
| | Sex * Day | 1, 13 | .034 | .001* | .506 |
| Conditioning Day 7-12 | Sex | 1, 13 | .544 | .474 | .040 |
| | Sex * Cue | 1, 13 | .271 | .611 | .020 |
| | Sex * Day | 5,65 | .110 | .990 | .008 |
| | Sex * Day * Cue | 5,65 | .606 | .696 | .045 |
| Blocking | Sex | 1,13 | .348 | .565 | .026 |
| | Sex * Cue | 2,26 | .977 | .390 | .070 |
| | Sex * Day | 3,39 | .492 | .690 | .036 |
| | Sex * Day * Cue | 6, 78 | 1.538 | .177 | .106 |
| Test | Sex | 1, 13 | 0.16 | .901 | .001 |
| | Sex * Cue | 1, 13 | 0.54 | .947 | .004 |
| **Experiment 2** $n_m = 24$; $n_f = 20$ | **Variable** | **df** | **F** | ***p*** | $\eta^2$ |
| Conditioning Day 1-6 | Sex | 1, 40 | .691 | .411 | .017 |
| | Sex * Group | 1, 40 | 2.522 | .120 | .059 |
| | Sex * Cue | 5, 200 | .665 | .650 | .016 |
| | Sex * Group * Cue | 5, 200 | .741 | .594 | .018 |
| Conditioning Day 7-12 | Sex | 1, 40 | .077 | .782 | .002 |
| | Sex * Group | 1, 40 | 1.573 | .217 | .038 |
| | Sex * Cue | 1, 40 | 2.112 | .154 | .050 |
| | Sex * Day | 5, 200 | .598 | .702 | .015 |
| | Sex * Group * Cue | 1, 40 | 1.024 | .318 | .025 |
| | Sex * Group * Day | 5, 200 | 1.165 | .328 | .028 |
| | Sex * Cue * Day | 5, 200 | .827 | .532 | .020 |
| | Sex * Group * Cue * Day | 5, 200 | .301 | .912 | .007 |
| Blocking | Sex | 1, 40 | .058 | .811 | .001 |

| | | | | | |
|---|---|---|---|---|---|
| | Sex * Group | 1, 40 | .052 | .822 | .001 |
| | Sex * Cue | 1, 40 | >.001 | .984 | >.001 |
| | Sex * Day | 3, 120 | .533 | .661 | .013 |
| | Sex * Group * Day | 3, 120 | .136 | .939 | .003 |
| | Sex* Day * Cue | 3, 120 | .860 | .464 | .021 |
| | Sex * Group * Cue * Day | 3, 120 | .726 | .539 | .018 |
| Test | Sex | 1, 40 | .004 | .949 | >.001 |
| | Sex * Group | 1, 40 | .065 | .799 | .002 |
| | Sex * Cue | 1, 40 | 1.499 | .228 | .036 |
| | Sex * Cue * Group | 1, 40 | .180 | .674 | .004 |
| **Experiment 3** $n_m = 19$; $n_f = 17$ | **Variable** | **df** | **F** | ***p*** | $\eta^2$ |
| Conditioning Day 1-6 | Sex | 1, 28 | 1.112 | .301 | .038 |
| | Sex * Group | 3, 28 | 1.654 | .199 | .151 |
| | Sex * Day | 5, 140 | .816 | .540 | .028 |
| | Sex * Group * Day | 15, 140 | .949 | .512 | .092 |
| Conditioning Day 7-12 | Sex | 1, 28 | .906 | .349 | .031 |
| | Sex * Group | 3, 28 | 1.330 | .285 | .125 |
| | Sex * Cue | 1, 28 | .013 | .908 | >.001 |
| | Sex * Day | 5, 140 | 1.177 | .324 | .040 |
| | Sex * Group * Cue | 3, 28 | .121 | .947 | .013 |
| | Sex * Group * Day | 15, 140 | | | |
| | Sex * Day * Cue | 5, 140 | .651 | .828 | .065 |
| | Sex * Group * Day * Cue | 15, 140 | .615 | .859 | .062 |
| Conditioning Day 12 | Sex | 1, 28 | .025 | .876 | .001 |
| | Sex * Group | 3, 28 | .271 | .846 | .028 |
| | Sex * Cue | 1, 28 | .128 | .723 | .005 |
| | Sex * Group * Cue | 3, 28 | .591 | .626 | .060 |
| Blocking Day 13-16 | Sex | 1, 28 | .418 | .523 | .015 |
| | Sex * Group | 3, 28 | .240 | .868 | .025 |
| | Sex * Cue | 1, 28 | .041 | .841 | .001 |
| | Sex * Day | 3, 84 | 1.790 | .155 | .060 |
| | Sex * Group * Cue | 3, 28 | .351 | .788 | .036 |
| | Sex * Group * Day | 9, 84 | .279 | .979 | .029 |
| | Sex * Day * Cue | 9, 84 | .269 | .848 | .010 |
| | Sex* Day *Cue * Group | 9, 84 | .852 | .571 | .084 |
| Blocking Day 16 | Sex | 1, 28 | .035 | .853 | .001 |
| | Sex * Group | 3, 28 | .167 | .918 | .018 |
| | Sex * Cue | 1, 28 | .285 | .597 | .010 |
| | Sex * Group * Cue | 3, 28 | .402 | .753 | .041 |
| Test | Sex | 3, 28 | .163 | .689 | .006 |

| | | | | | |
|---|---|---|---|---|---|
| | Sex * Cue | 1, 28 | .296 | .591 | .010 |
| | Sex * Group | 3, 28 | 1.069 | .378 | .103 |
| | Sex * Cue * Group | 3, 28 | .639 | .596 | .064 |

Conditioning. TH-Cre rats were trained in a within-subjects blocking design. On Days 1-6 (not tethered) of conditioning, rats received 16 presentations of a 10s visual stimulus A paired with two sucrose rewards at 9s and 10s (A → reward). A mixed ANOVA revealed an increase in responding across days (Supplemental Fig 1A; $F_{Day}(5,70) = 14.364$, $p = .857$, $\eta^2 = .003$), a main effect of sex ($F_{Sex}(1,13) = 5.205$, $p = .040$, $\eta^2 = .286$), but no interaction ($F_{Day*Sex}(1,13) = .034$, $p < .0001$, $\eta^2 = .506$). This main effect of sex did not persist for the rest of the experiment. On Days 7-12, in addition to the A → reward trials, a non-reinforced visual stimulus B was introduced (B → nothing) where three presentations of B were randomly intermixed with eight presentations of A and this block was repeated twice (i.e., 16x A → reward; 6x B → nothing). A repeated ANOVA revealed an increase in responding across days ($F_{Day}(5,70) = 3.835$, $p = .004$, $\eta^2 = .215$) and a successful discrimination between the reinforced A and the non-reinforced B ($F_{Cue}(1,14) = 26.054$, $p < .0001$, $\eta^2 = .650$), and no interaction between days and cues ($F_{Cue\ x\ Day}(5,70) = .465$, $p = .801$, $\eta^2 = .032$).

Blocking. Blocking took place across days 13-16 and consisted of three audio-visual compounds (AX, AY, BZ) each paired with reward. Optical stimulation of VTA TH+ neurons occurred on AX → reward during reward delivery at 9s and 10s and AY → reward trials during the intertrial interval. Responding was high and remained stable for all compounds containing A (i.e., AX, AY), but increased across days for the compound containing B (i.e., BZ; Supplemental Fig 1A). A repeated measures ANOVA revealed no effect of days ($F_{Day}(3, 42) = 1.064$, $p = .375$, $\eta^2 = .071$), an effect of cues ($F_{Cue}(2,28) = 3.734$, $p = .037$, $\eta^2 = .211$) but no interaction ($F_{Cue\ x\ Day}(6,84) = .652$, $p = .689$, $\eta^2 = .044$). There were no differences between the compounds by the end of blocking (AX vs. AY: $M_{diff} = 2.088$, $SE = 1.115$, $p = .247$, 95% CI [-.943, 5.118]; AX vs. BZ $M_{diff} = 3.362$, $SE = 1.382$, $p = .087$, 95% CI [-.393, 7.117]); AY vs BZ: $M_{diff} = 1.275$, $SE = 1.216$, $p = .937$, 95% CI [-2.030, 4.579]).

Test. On test (Supplemental Fig 1A), non-reinforced presentations of X, Y and Z revealed lower level of responding to the blocked cue Y and unblocking to X relative to the control cue Z. A repeated measures ANOVA confirmed these observations ($F_{Cue}(2,28) = 6.070$, $p = .006$, $\eta^2 = .302$). Post-hoc analyses revealed that responding to the blocked cue Y was lower than to the control cue Z ($M_{diff} = -10.320$, $SE = 3.307$, $p = .023$, 95% CI [-1.333, -19.307) while responding to the unblocking cue X was greater than the blocked cue Y ($M_{diff} = 8.011$, $SE = 2.550$, $p = .022$, 95% CI [1.080, 14.942). These data show that stimulation of VTA TH+ neurons at time of expected reward unblocks learning, and furthermore, that it does so to the level of the controls (i.e., responding to X did not differ from Z, ($M_{diff} = -2.309$, $SE = 3.400$, $p = 1.00$, 95% CI [-11.550, 6.932]). That is, it yielded complete unblocking.

**Experiment 2: A value upshift halfway through conditioning does not disrupt the blocking effect.**

In the experiment examining the effect of optical stimulation of VTA DA neurons during reward delivery in both phases of a blocking design, stimulation began half way through the conditioning session. To be sure that increasing the value of the reward half way through conditioning did not lead to unblocking, we conducted a behavioural experiment in which one group of rats received an upshift in reward from conditioning to blocking (Group Unblock) whereas another group received the higher reward during conditioning and blocking (Group Block). The data confirmed that a change in US intensity from conditioning to blocking led to unblocking, whereas delivering the same reward albeit large reward during conditioning and blocking led to blocking.

Conditioning. This experiment was done as a precursor to Experiment 3 in order to determine whether increasing the value of the reward paired with A half way through Phase 1, as would be done using optical stimulation in Experiment 3, would yield blocking. Two groups of WT rats were trained in a within-subjects blocking design similar to that described previously (Supplemental Fig 3A). Group Unblock received A → 1x reward (i.e., one pellet) throughout conditioning where as Group Block received A → 1x reward for the first 6 days of conditioning and A → 3x reward (i.e., three pellets) for the rest of conditioning (Days 7-12). A mixed ANOVA on Days 1-6 revealed an increase in responding across days (Supplemental Fig 3B; $F_{Day}(5, 210) = 57.002$, $p < .0001$, $\eta^2 = .578$), no effect of group ($F_{Group}(1,42) = .631$, $p = .432$, $\eta^2 = .015$) and no interaction ($F_{Day*Group}(5, 210) = .483$, $p = .789$, $\eta^2 = .011$). On days 7-12, A → reward trials were accompanied by B→ nothing trials. A mixed ANOVA confirmed an increase in responding across days 7-12 ($F_{Day}(5,210) = 6.578$, $p <.0001$, $\eta^2 = .135$) with no main effect of group ($F_{Group}(1, 42) = .004$, $p = .952$, $\eta^2 < .001$), nor group interactions (max $F(1, 42) = 2.082$, $p = .156$, $\eta^2 = .047$). There was a main effect of cue ($F_{Cue}(1,10) = 139.803$, $p < .0001$, $\eta^2 = .769$) and a cue by day interaction ($F_{Cue * Day}(5, 210) = 8.784$, $p < .0001$, $\eta^2 = .173$), such that rats learned to discriminate the A and B across training days and were able to discriminate the cues on day 12 ($M_{diff} = 24.782$, $SE = 2.515$, $p < .0001$, 95% CI [19.706, 29.858]).

Blocking. During Blocking, responding to AX remained high whereas that to BY increased across days (Supplemental Fig 3B). A mixed ANOVA showed an increase in responding across days ($F_{Day}(3, 126) = .446$, $p <.0001$, $\eta^2 = .011$) and no main effect of group ($F_{Group}(1,42) = .198$, $p = .659$, $\eta^2 = .005$) nor group interactions (max $F(1, 42) = 2.164$, $p = .149$, $\eta^2 = .049$). There was a main effect of compound ($F_{Cue}(1,42) = 9.674$, $p = .003$, $\eta^2 = .187$) and an interaction between the compounds and training days ($F_{Cue\ x\ Day}(3, 126) = 2.752$, $p = .045$, $\eta^2 = .061$) which suggest the compound effect disappeared by the end of blocking. Post hocs confirmed that responding differed between AX and BY on Day 1(Group Block: $M_{diff} = 3.529$, $SE = 1,397$, $p = .015$, 95% CI [.709, 6.349]; Group Unblock: $M_{diff} = 3.805$, $SE = 1.397$, $p = .009$, 95% CI [-6.624, -.985]) but not by the end of blocking on Day 4 (Group Block: $M_{diff} = .425$, $SE =$

1.398, $p$ = .763, 95% CI [-3.247, 2.397]; Group Unblock: $M_{diff}$= 1.655, $SE$ = 1.398, $p$ = .243, 95% CI [-1.169, 4.476]).

Test. Test to X and Y showed blocking in Group Block evident in lower levels of responding to X compared to Y and a loss of this difference (i.e., unblocking) in Group Unblock (Supplemental Fig 3B). A mixed ANOVA confirmed these observations, showing no effect of group ($F_{Group}(1,42)$= .287, $p$ = .595, $\eta^2$ = .007), no effect of cue ($F_{Cue}(1,42)$= 2.100, $p$ = .155, $\eta^2$ = .048) with a group by cue interaction ($F_{Cue\ x\ Group}(2,42)$= 5.814, $p$ = .020, $\eta^2$ = .122). Post-hoc comparisons revealed that responding to X was lower compared to Y in the Group Block ($M_{diff}$= 11.152, $SE$ = 4.085, $p$ = .009, 95% CI [2.907, 19.396]) but not in Group Unblock ($M_{diff}$= -2.779, $SE$ = 4.085, $p$ = .500, 95% CI [-11.023, 5.465]).

**Experiment 3: Optogenetic stimulation of VTA DA transients mimics prediction error not value.**

Conditioning. Four groups of TH-cre rats (AX $n$ = 10; eYFP $n$ = 8; ITI $n$ = 10; BY $n$ = 8) were trained in a within-subjects blocking design similar to that described previously (Figure 2A). The difference between each of the four groups consisted of the timing of optical stimulation of TH+ neurons. Group AX received stimulation during reward delivery on A→reward (Conditioning) and AX→reward (Blocking) trials to determine if stimulation serves to add value to the reward or induces a prediction error. To control for light artifacts, temporal specificity of stimulation and value to an unexpected reward, three control groups were generated. Group eYFP (lacking the opsin in TH+ neurons) received identical optical stimulation to Group AX; Group ITI was stimulated during the ITI in both phases of learning (Conditioning, Blocking); Group BY received stimulation during the ITI in Conditioning and reward delivery in Blocking control, that is, BY reward trials.

Responding to the reinforced cue A increased across the first six days for all groups (Supplemental Fig 2A). A mixed ANOVA confirmed an effect of days ($F_{Day}(5, 160)$ = 72.516, $p$ < . 0001, $\eta^2$ = .694) and no effect of group ($F_{Group}(3, 32)$ = .647, $p$ = .591, $\eta^2$ = .057). There was a day by group interaction ($F_{Day * Group}(15, 160)$ = 1.968, $p$ = .021, $\eta^2$ = .156). On days 7-12, non-reinforced B trials were added and optical stimulation began. Discrimination between A and B was successful for all the groups. A mixed ANOVA confirmed an effect of days ($F_{Day}(5,160)$ 4.610, $p$ = .001, $\eta^2$ = .126), an effect of cue ($F_{Cue}(1,32)$ = 118.270, $p$ < .001, $\eta^2$ = .787) and no effect of group ($F_{Group}(3, 32)$ = .803, $p$ = .502, $\eta^2$ = .070). There was a cue by day interaction ($F_{Cue\ x\ Day}(5,160)$ = 4.328, $p$ = .001, $\eta^2$ = .119), showing that the discrimination was acquired across days. The rate of acquiring the discrimination differed between groups ($F_{Cue\ x\ Day\ x\ Group}$ (5,160) = 1.777, $p$ = .042, $\eta^2$ = .143) and the magnitude of the difference in responding to A and B differed between groups ($F_{Cue\ x\ Group}(3,32)$ = 4.838, $p$ = .007, $\eta^2$ = .312). Importantly, on the final day of conditioning (day 12), all groups were able to discriminate A and B ($F_{Cue}$ (3,32) = 85.336, $p$ < .001, $\eta^2$ = .727) with no group differences ($F_{Group}$ (3,32) = 1.040, $p$ = .388, $\eta^2$ = .089; $F_{Cue\ x\ Group}$ (3,32) = 2.844, $p$ = .053, $\eta^2$ = .211).

Blocking. During blocking, A was presented with X, and B with Y to generate a blocking and control condition, respectively. Groups AX and eYFP received optical stimulation occurred during reward delivery for AX trials, Group ITI received optical stimulation during the ITI following AX trials, Group BY received optical stimulation at the time of reward following BY. As expected, responding to AX was high at the start of blocking whereas that to BY increased across days for all groups, with no differences between the compounds by the end of blocking (Supplemental Fig 2A). This was confirmed by the statistical analyses. A mixed ANOVA revealed an effect of cue ($F_{Cue}(1,32) = 9.077$, $p = .005$, $\eta^2 = .221$), an effect of days ($F_{Day}(3,96) = 36.794$, $p < .001$, $\eta^2 = .535$), no effect of group ($F_{Group}(3, 32) = .265$, $p = .850$, $\eta^2 = .024$) and a cue by day interaction ($F_{Cue * Day}(3, 96) = 2.877$, $p = .040$, $\eta^2 = .082$). While the rate of the increase in responding to BY differed by group ($F_{Cue * Group * Day}(9, 96) = 2.215$, $p = .027$, $\eta^2 = .172$), there were no group differences overall in responding to the compounds ($F_{Cue * Group}(3, 32) = 2.193$, $p = .108$, $\eta^2 = .171$). Importantly, there were no group differences in responding to the compounds on the final day of blocking (Day 16: $F_{Cue}(1, 32) = 1.266$, $p = .269$, $\eta^2 = .038$; $F_{Cue * Group}(3, 32) = .642$, $p = .594$, $\eta^2 = .057$).

Test. Test to X and Y revealed blocking across all of the controls groups (eYFP, ITI, BY) except Group AX, which showed unblocking to X (Supplemental Fig 2A). A mixed ANOVA confirmed this observation, showing no effect of group ($F_{Group}(3,32) = .071$, $p = .975$, $\eta^2 = .007$), an effect of cues ($F_{Cue}(1,32) = 28.098$, $p < .001$, $\eta^2 = .468$) and an interaction ($F_{Cue\ x\ Group}(3,32) = 3.757$, $p = .020$, $\eta^2 = .260$). Post-hoc comparisons confirmed that responding to X was less than to Y for Group eYFP ($M_{diff}$= -15.912, *SE* = 4.542, $p = .001$, 95% CI [-6.662, -25.163]), Group ITI ($M_{diff}$= -10.000, *SE* = 4.040, $p = .019$, 95% CI -18.230, -1.771]) and Group BY ($M_{diff}$= -19.087, *SE* = 4.57, $p < .001$, 95% CI [-28.289, -9.886]), but not for Group AX ($M_{diff}$= -.430, *SE* = 4.062, $p = .916$, 95% CI [-8.704, 7.844]), thus confirming that optical stimulation of VTA DA neurons across both conditioning and blocking phases unblocked learning.

#### METHODS

##### Subjects

The optogenetic experiments consisted of 54 (Experiment 1: 18, 8 males, 10 females; arrival weight 250-350g; Experiment 3: 44 males, females; weight 250-420g) transgenic rats expressing Cre recombinase under the control of the tyrosine hydroxylase (TH) promoter on a Long Evans background (TH:Cre rats). Due to incorrect fiber placements following histological analyses, the final number of rats included in the analyses was 15 (7 males, 8 females) in Experiment 1 and 36 (18 males, 18 females) in Experiment 3. Experiment 2 consisted of 44 Long Evan wild type rats (24 males, 20 females: weight 250-350g). The rats for Experiment 1 were obtained from (the National Institute on Drug Abuse, MD, USA). All other rats were bred in-house in the Animal Care Facility at Concordia University. Concordia University, QC, CA). Rats were pair-housed in a temperature (21°C) and humidity (44%) controlled colony room on a 12 h reverse light/dark cycle (lights on at 9 AM; lights off at 9 PM) in clear standard cages (44.5cm x 25.8 cm x 21.7 cm) lined with a mix of beta chip and corncob bedding (Teklad, Envigo). Water and food (18% protein, 4.5% fat; Charles River) were available *ad libitum* prior to surgery and during recovery. During the experiment, access to food was restricted to approximately 10g/female/day and 13g/male/day to achieve and maintain 85% of their recovery body weight. All experimental procedures were in accordance with the Canadian Council on Animal Care and the Concordia University Animal Care Committee.

**Housing**

Rats were pair-housed in a temperature (21°C) and humidity (44%) controlled colony room on a 12 h reverse light/dark cycle (lights on at 9 AM; lights off at 9 PM) in clear standard cages (44.5cm x 25.8 cm x 21.7 cm) lined with a mix of beta chip and corncob bedding (Teklad, Envigo). Water and food (18% protein, 4.5% fat; Charles River) were available *ad libitum* prior to surgery and during recovery. During the experiment, access to food was restricted to approximately 10g/female/day and 13g/male/day to achieve and maintain 85% of their recovery body weight.

**Behavioral Procedures**

**Experiment 2.**

All subjects were handled by the experimenter for five days prior to the start of the experiment. Behavioral training was conducted during the dark cycle. Sessions began with a 2 min adaptation period. Each reinforced conditioning trial consisted of a 10s cue (or compound) followed by pellet delivery at 9 s and 10 s. Non-reinforced trials consisted of cue alone presentations. The intertrial interval lasted on average 6 min (range: 4-8 min). Rats remained in the conditioning chamber for 2 min following the final trial. The walls, ceiling, floor and stainless-steel tray were cleaned with a 10 % acetic acid solution before each session.

<u>Phase 0 Magazine Training.</u> Rats were exposed to sucrose pellets (20 pellets/rat) in their home cages 24 h prior to magazine training. Each magazine training session lasted 40 min during which one sucrose pellet was delivered into the food port every 60 s.

<u>Phase 1 Conditioning.</u> Conditioning took place across 12 days. On days 1-6, rats received 16 presentations of a 10 s stimulus (A) paired with two sucrose pellets. On days 7-12, six non-reinforced presentations of visual cue (B) were intermixed with the 16 reinforced trials of A in mini blocks of two such that B was presented randomly three times for every eight presentations of A. In experiments 1 and 3, rats were habituated to the background no ise from the laser equipment one day prior to use.

<u>Phase 2 Blocking</u>. Each compound conditioning session began with eight reminder trials of A and B presented in the following order AABBAABB and counterbalanced for initial cue presentation across days. Following reminder trials, compounds were presented six times each in a pseudo-randomized order. For experiment 1, this order was XXYYZZ ZZYYXX YYZZXX. For experiment 2 and 3 the order was XYYXYXXYXYXY for day 1 and 3 and XYXYXXYXYYXY for days 2 and 4.

<u>Test.</u> To determine the level of conditioned approach acquired by the auditory stimuli, rats received three non-reinforced presentations of each cue. In experiment 1 the order was: XYZ ZXY YZX, with two reinforced presentations of A after each set of 3 cues. In experiment 2 and 3 the order was XYYXYXY. All pseudo-randomized cues were counterbalanced for the initial stimulus presentation.

### Supplemental Figure 1.

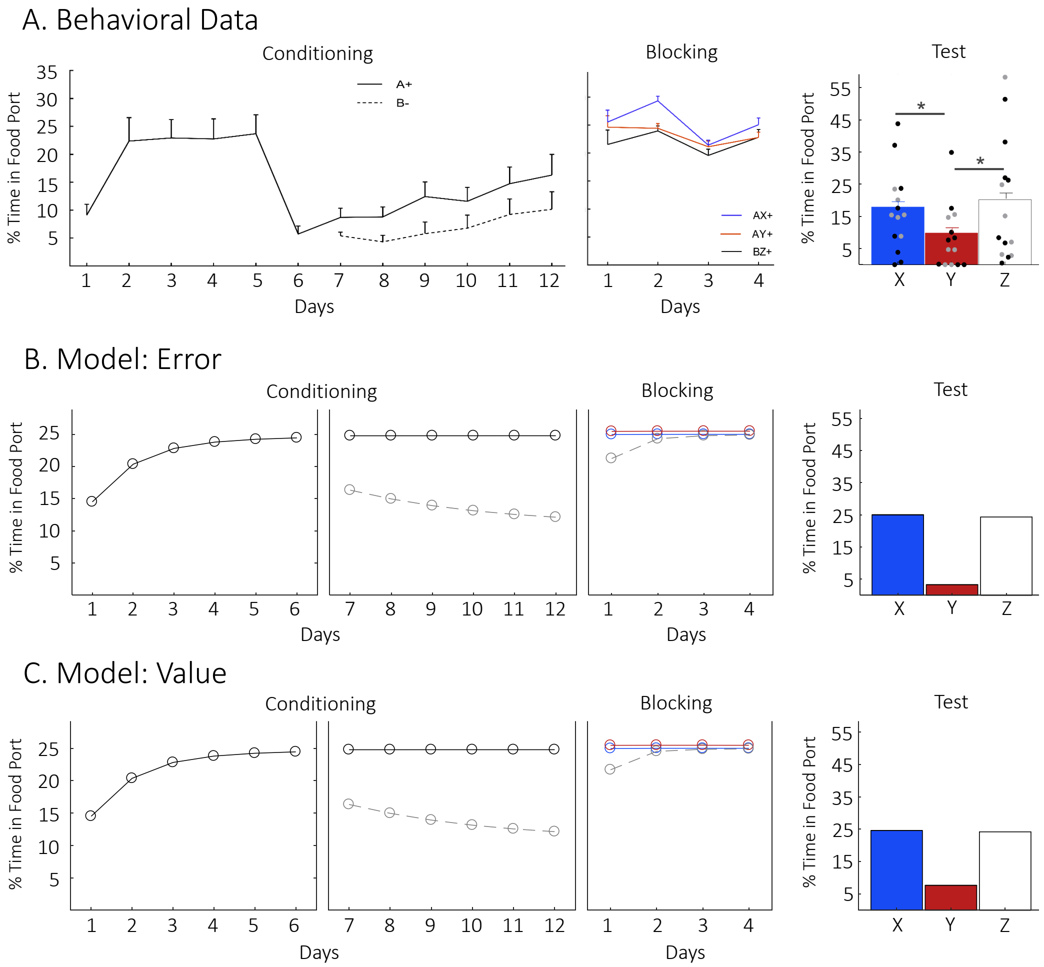


**Supplemental Figure 1. Optogenetic stimulation of VTA DA transients promotes learning in blocking**.
A) Behavioural data (mean + s.e.m., *n* = 15 rats; males in gray, females in black) obtained during the conditioning, blocking and test; B) Predicted results based on computational modeling where VTA DA stimulation acts as a prediction error (Model: Error) or C) as a rewarding event (Value Model). Note the output of the classic temporal difference reinforcement learning model was converted from *V* to CR to better reflect the behavioral output actually measured in our experiments (see <u>Methods</u> for details).

#### Supplemental Figure 2.

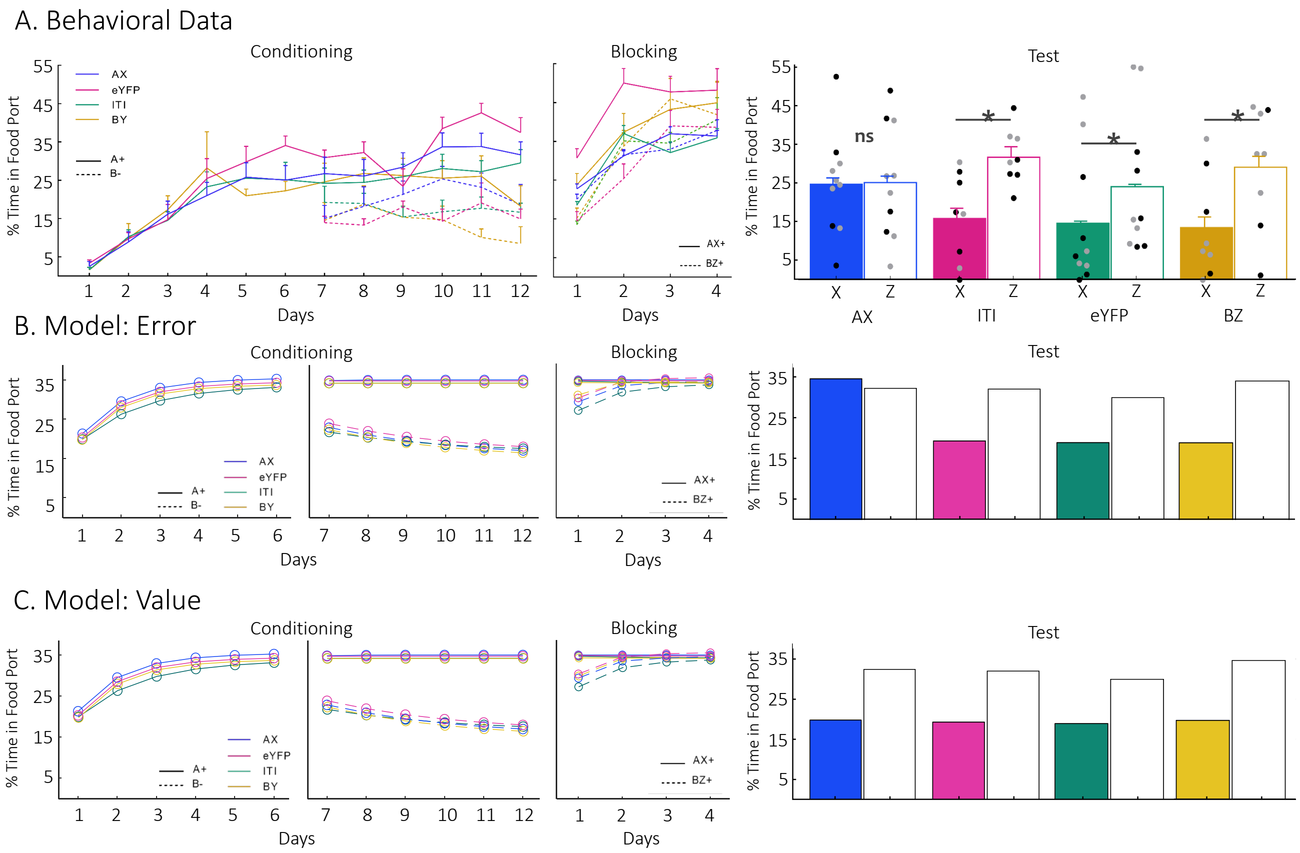


**Supplemental Figure 2. Optogenetic stimulation of VTA DA transients generate a prediction error, not a rewarding event.** A) Behavioural data (mean + s.e.m., *n* = 44 rats; males in gray, females in black) obtained during the conditioning, blocking and test; B) Predicted results based on computational modeling where VTA DA stimulation acts as a prediction error (Model: Error) or C) as a rewarding event (Value Model). Note the output of the classic temporal difference reinforcement learning model was converted from *V* to CR to better reflect the behavioral output actually measured in our experiments (see Methods for details).

#### Supplemental Figure 3.

##### A. Design

| Group | Conditioning | | Blocking | Test |
|---|---|---|---|---|
| Block | A+ | A+++ | AX+++ | X |
| | | B- | BZ+++ | Z |
| Unblock | A+ | A+ | AX+++ | X |
| | | B- | BZ+++ | Z |

##### B. Behavioral Data

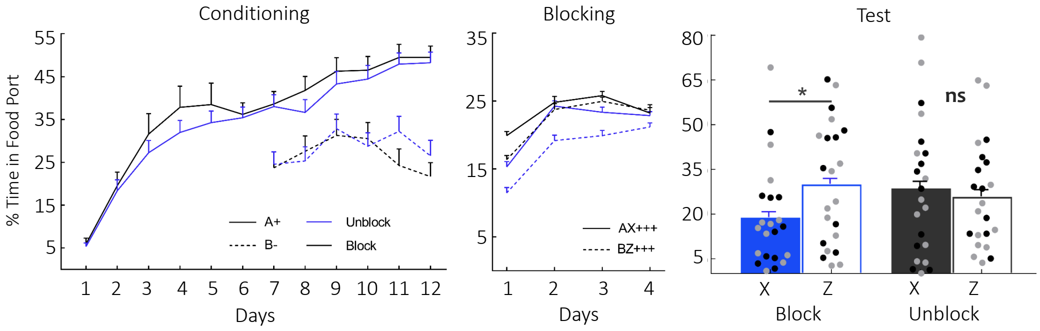


**Supplemental Figure 3. A value upshift halfway through conditioning does not disrupt the blocking effect. A)** Behavioral design. **B)** Behavioral results (mean + s.e.m., *n* = 44 rats; males in gray, females in black) on test confirm a cue by group interaction (ANOVA: $F_{Cue\ x\ Group}(2,42)$= 5.814, $p$ = .020, $\eta^2$ = .122) where responding to X was lower than to the control cue Z for group Block ($M_{diff}$= 11.152, *SE* = 4.085, $p$ = .009, 95% CI [2.907, 19.396]) but not in group Unblock ($M_{diff}$ = -2.779, *SE* = 4.085, $p$ = .500, 95% CI [-11.023, 5.465]).
